# Value-Added: Importance of Incorporating Menstrual Cycle Phases to Clarify Sex-Related Differences in Force Steadiness

**DOI:** 10.1101/2025.08.27.672658

**Authors:** Cori A. Calkins, Derek C. Chin, Kathryn M. Crosby, Elijah M. K. Haynes, Jennifer M. Jakobi

## Abstract

**Aim:** This study aimed to determine whether female elbow flexion force steadiness varies across the menses, follicular and luteal phases of the menstrual cycle.

**Methods:** To control for repeated testing effects unrelated to hormonal fluctuations across the menstrual cycle, a comparison group of males completed the same protocol over three equally spaced testing sessions to the females. Maximal voluntary contractions and force steadiness were assessed in the neutral and pronated forearm positions. Elbow flexion force tracking tasks were performed at 2.5, 5, 10, 25, 50 and 75% maximal voluntary contraction in both forearm positions, and force steadiness was quantified as the coefficient of variation of force.

**Results:** Males were stronger than females (p<0.001), and maximal voluntary contractions did not differ between menstrual phases (p>0.14) or between days in males (p>0.56) in both forearm positions. There was no difference in coefficient of variation of force across sessions for the males (p<0.36). The coefficient of variation of force for all submaximal forces was significantly greater during the luteal phase compared to menses (Neutral p=0.02; Pronated p<0.05) but not the follicular phase (Neutral p=0.71; Pronated p=0.10). The coefficient of variation of force during the luteal phase in females was higher than males in both positions (p<0.02).

**Conclusion:** These findings support previous observations that females are less steady than males for isometric steady contractions; however, this study identifies luteal specific phase effects. This underscores the importance of accounting for menstrual cycle phase when conducting sex-related comparisons.

## Introduction

Males typically produce more forceful and steadier voluntary contractions compared to females (Haynes et al., 2020, Brown et al., 2010). These sex-based differences in force output and modulation could be in part due to menstrual cycle phase whereby cyclical fluctuations of sex hormones occur, which may influence the ability to maintain a steady contraction (force steadiness) as estrogen and progesterone receptors have been identified on skeletal muscle tissue and throughout the nervous system (Gargiulo-Monachelli et al., 2014; Lemoine et al., 2003). Although sex-based differences in force steadiness have been investigated, far less attention has been given to determining the influence of menstrual phase on force steadiness (Jakobi et al., 2018; Tenan et al., 2016).

One potential reason for this research gap is the perceived complexity of studying hormonal contributors that fluctuate over time, such as the menstrual cycle in females (Allen et al., 2016). While there are both perceived, and practical challenges such as resource limitations for sample storage, analysis or associated costs of measuring hormones it is clear that there are noticeable gaps in research studies on females (Inglis & Cabral, 2025). Nevertheless, studies that assess force steadiness across the menstrual cycle, even in the absence of direct hormone measurements, are valuable, as they can offer meaningful insight into phase-related trends, and this approach accounts for the fact that females function on a daily basis across phases rather than on known and fixed hormonal levels.

Force steadiness is influenced by various biomechanical factors. For example position, where a pronated forearm position increases the amplitude of force fluctuations compared to a neutral position (Brown et al., 2010; Smart et al., 2018). Steadiness also varies with contraction intensity, with the steadiest forces typically occurring at moderate force levels and least steady at low and high force levels (Brown et al., 2010). Accordingly, the contribution of the motor unit pool is also known to vary between low and high forces (Dideriksen et al. 2012; Farina & Negro 2015), and thus rigorous evaluation of elbow flexion force steadiness across the menstrual cycle should account for both forearm position and contraction intensity.

Despite growing interest in sex-related differences in force control and modulation (Lulic-Kuryllo & Inglis, 2022; Jakobi et al., 2018) it remains unknown whether force steadiness varies across the menstrual cycle. This study aimed to determine the influence of menstrual phase on force steadiness in eumenorrheic females not using hormonal contraceptives. A secondary aim was to assess sex-based differences in force steadiness by including males over comparable time intervals. It was hypothesized that force steadiness would vary by menstrual phase, with the luteal phase exhibiting the highest CV of force, and that sex-related differences would be most evident in this phase.

## Materials and Methods

### Eligibility and Recruitment

Young adults between the ages of 19 to 35 years were recruited to participate in this study. Individuals were excluded if they had a cardio-metabolic or neurological condition, participated in elite levels of upper-body resistance training, undergone surgery or sustained a severe injury to the neck, shoulder or arms, or had taken hormonal contraceptives in the previous six months. The study was approved by the University of British Columbia Behavioural Research Ethics Board (H16-00948-A010).

### Experimental Set-up

Participants were seated upright in a custom-built dynamometer chair that was adjusted such that their hips and knees were flexed at 90°, and their feet were planted on a flat surface.

The elbow joint of the dominant arm was flexed at 110° and supported beneath the olecranon. The shoulder was slightly abducted at an approximate angle of 30°. The participant’s dominant hand firmly gripped the manipulandum in the pronated (palm down) or neutral (thumb up) forearm positions. A linearly calibrated force transducer (MLP-150, 68 kg, 266 V sensitivity, Transducer Techniques, Temecula, CA, USA) fixed between the manipulandum and chair arm was used to measure elbow flexion force. The force signal was collected via a bridge amplifier (7.5 V excitation, 266 V sensitivity, Coulbourn Electronics, Allentown, PA, USA) and sampled at 1024 Hz using a Power 1401 analog-to-digital converter (Cambridge Electronic Design, Cambridge, England). Elbow flexion force was displayed on a 52 cm monitor aligned with the line of sight one metre in front of the participants. Force signals were stored locally and analyzed offline (MATLAB, Version 2021A, Massachusetts US).

### Experimental Protocol

Each participant executed the same protocol over three separate lab visits. For females, the three visits were randomized via menstrual cycle phase: menses (days 1-5), follicular (days 6-12) and luteal (day 16 to end of cycle (∼day 28)). To distinguish if there are any repeated testing session effects on CV of force unrelated to menstrual cycle influence, males also completed three testing sessions over a month. Each participant’s three visits began within one hour of the same time of day to mitigate variability due to diurnal menstrual cycle effects (Piirainen et al., 2023). For females, the order of testing was randomized between phases. During each visit, participants performed three 5-7 second MVCs in the neutral or pronated forearm position with the initial position randomized between participants. Participants rested between each MVC. The highest MVC in each position was used to normalize target force (% MVC).

Participants then performed two sets of submaximal tracking tasks at six force levels (2.5, 5, 10, 25, 50, and 75%) in a within-block randomized design. The blocks reflected the neutral and pronated forearm positions, and the block order was also randomized between participants and across days. During these submaximal tracking tasks, participants increased elbow flexion force over a 3-second ramp to the target force level. Participants were instructed and verbally encouraged throughout each task to sustain elbow flexion force at the target for 10 seconds and then, over 3 seconds, lower force output until full relaxation.

### Data Analysis

MVC force was analyzed during each experimental visit (Spike2, version 7.0, Cambridge, UK). The force signals were low pass filtered at 30 Hz and CV of force (standard deviation expressed as a percentage of mean force) during the middle 8 seconds of the plateau phase was analyzed offline (MATLAB, version 2021A, Massachusetts US).

### Statistical Analysis

Linear mixed effects (LME) models were conducted using the software R (R version 4.3.2, The R Foundation for Statistical Computing). Prior to all statistical analyses the extreme outliers were detected with box plots for each dependent variable, and values outside the 3rd quartile + 3 × interquartile range and 1st quartile – 3 × interquartile range were removed from analysis. Q-Q plots and histograms were used to access the assumptions of linearity, normality and homoscedasticity of the residuals. All residuals were found to be normal and linear.

To assess CV of force for the females between the three phases (menses, follicular, and luteal) at low and high force levels LME models were fit using package lme4 (Bates et al., 2015) with Satterthwaite’s method lmerTest (Kuznetsova et al., 2017). Cohen’s *d* values between force levels were calculated with effects size (Torchiano, 2020). Phase (i.e., menses, follicular and luteal) and force level (low (2.5, 5, 10, 25% MVC) and high (50 and 75%) were coded as fixed effects with participant as a random intercept. Each phase was further compared at each force level (2.5, 5, 10, 25, 50 and 75%) for each position with separate LME models. Phase was treatment-contrast coded and force level was sum-contrast coded. Example model: CV of Force ∼ Phase*Force Level + (1|Particpant).

To access CV of force across testing sessions in males LME model was fit with day (1, 2 and 3) as a fixed effect and participant as a random intercept. The *a priori* hypothesis was males would not differ between days, and subsequently when statistically proven the first day for the males was used in the sex-based LME model to assess CV of force in males (day 1) to the three phases of the menstrual cycle.

## Results

### Phase Effects

Twelve females (21.8 ± 2.0 years, 166.8 ± 6.4 cm, 63.8 ± 11.0 kg) and twelve males (22.8 ± 2.2 years, 179.9 ± 5.3 cm, 80.3 ±14.4 kg) participated in the study. In the neutral (p=0.02) and pronated (p<0.05) position there was an effect of menstrual phase, as CV of force in the luteal phase was greater than menses. There was no significant difference in CV of force between the follicular phase and menses (Neutral p=0.06; Pronated p=0.11) and follicular with luteal (Neutral p=0.71; Pronated p=0.10) (Figure 1). There was a significant effect of force level whereby the high forces (50% and 75% MVC) had lower CV of force than the low forces (2.5%, 5%, 10% and 25% MVC) for the neutral (p=0.02) and pronated (p<0.001) position (Figure 2).

**Figure 1.**
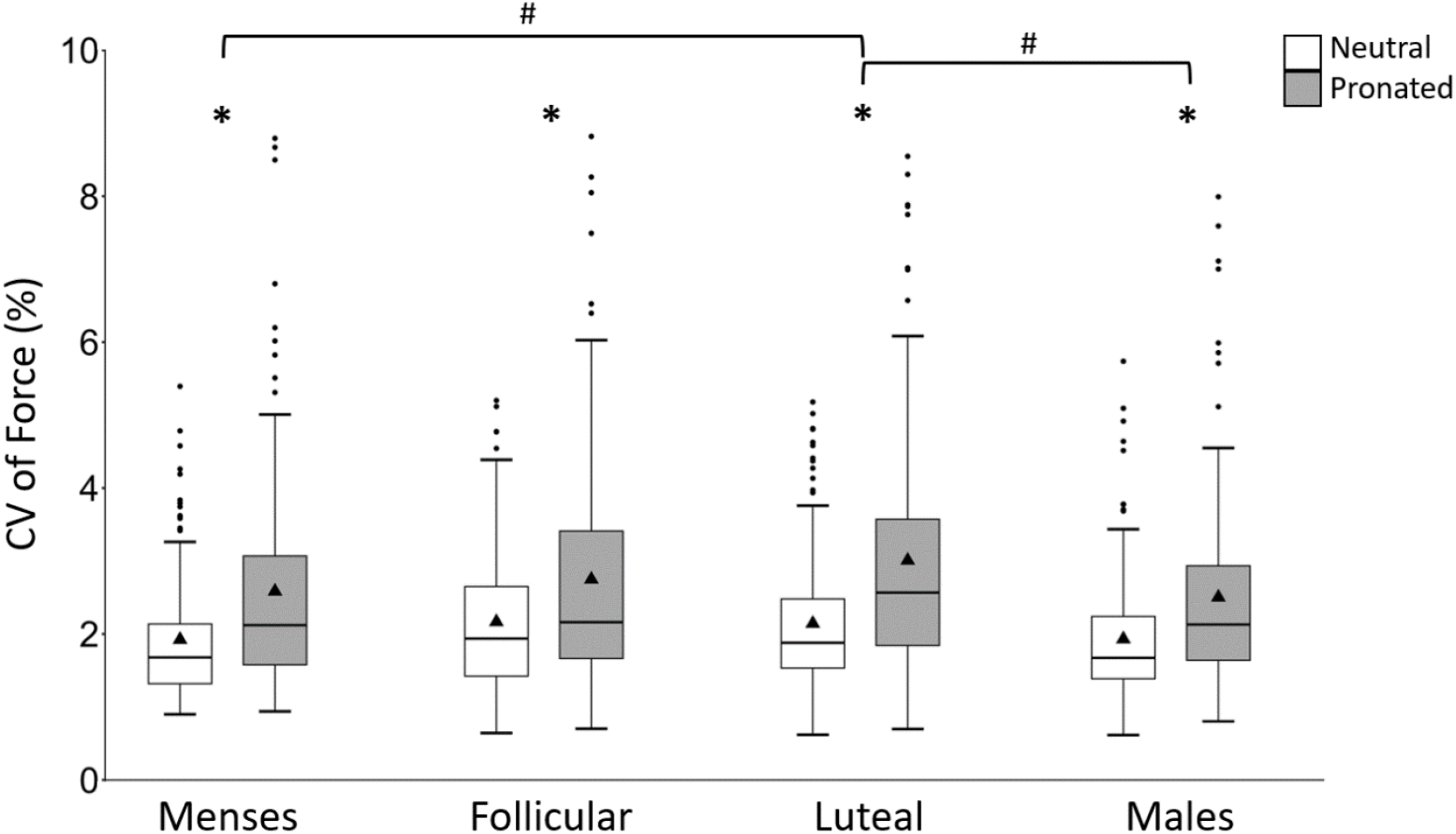
CV of force (%) of females across the menstrual cycle and males day 1. The filled in triangle denotes the mean. * Indicates a significant difference between neutral and pronated positions. # Indicates the luteal phase is significantly higher than menses, and the luteal phase is higher than males in the neutral and pronated positions p<0.05.

**Figure 2.**
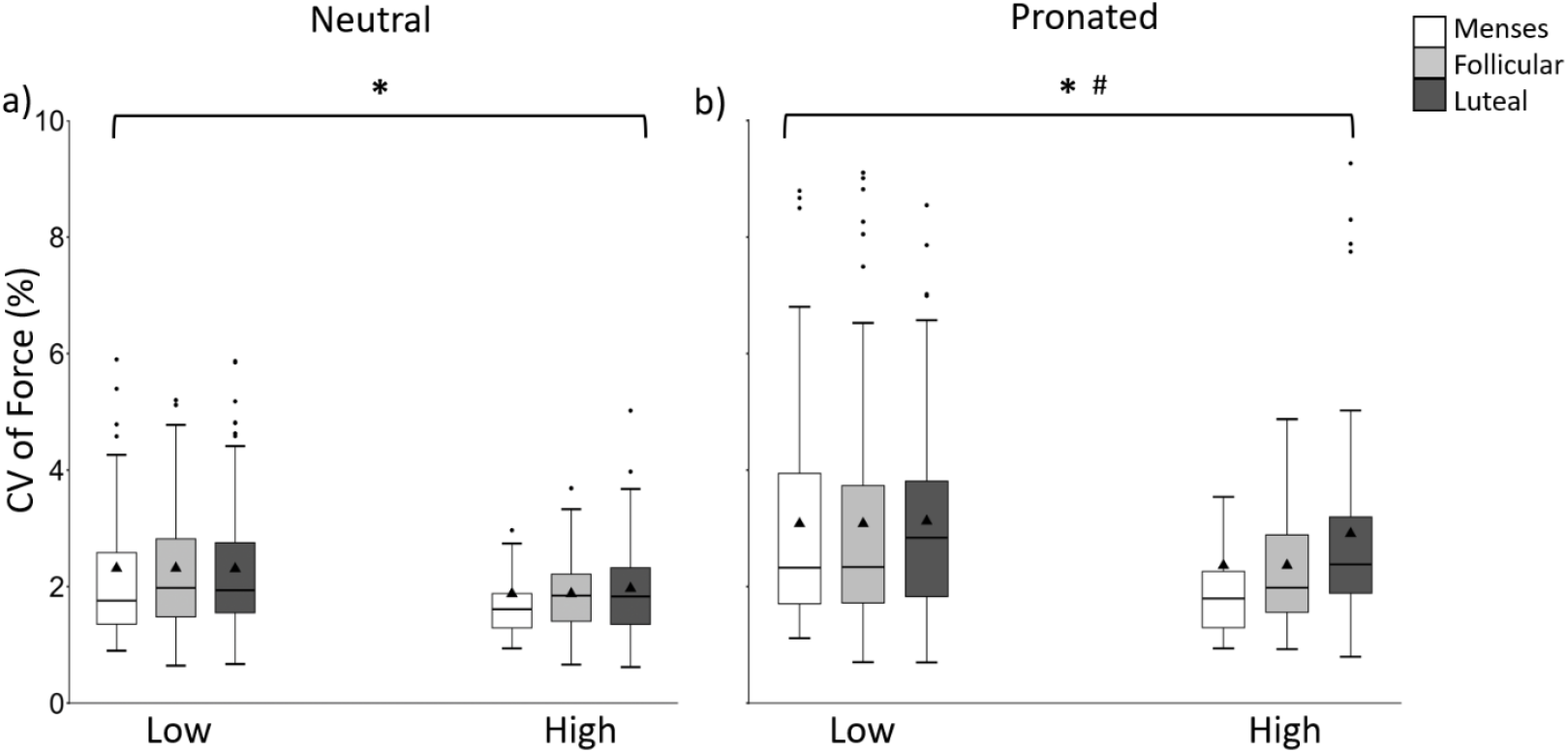
CV of force between low (2.5, 5, 10 and 25% MVC) and high (50, 75% MVC) force levels across the menstrual cycle in the neutral (a) and pronated (b) forearm positions. The filled triangles denote the mean. * Indicates a significant difference between low and high force levels. # Indicates a phase by force interaction where the difference between menses and the luteal phase between low and high force levels is significantly different. p<0.05.

**Figure 3.**
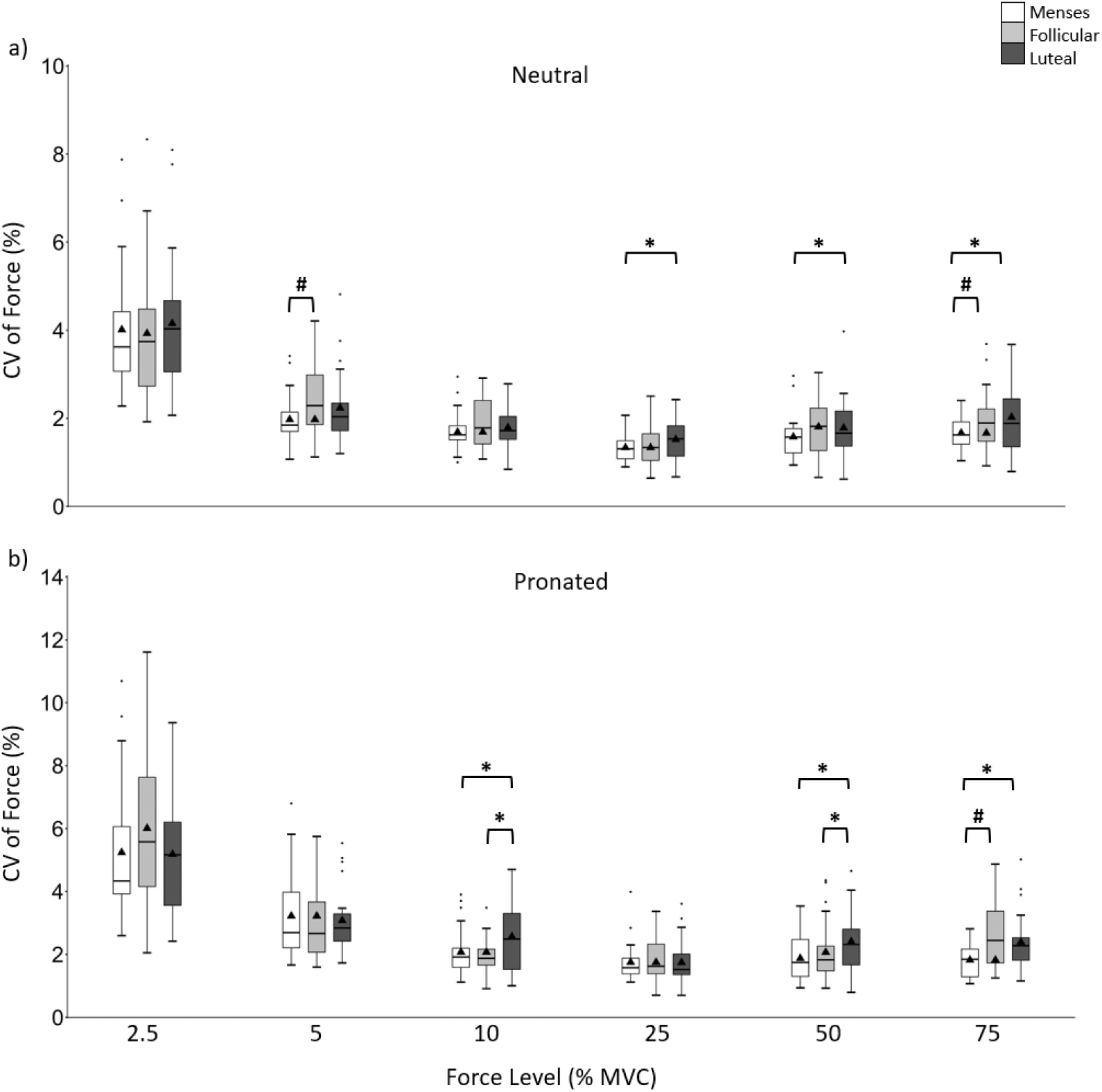
CV of force during isometric elbow flexion at each force level (i.e. 2.5, 5, 10, 25, 50 and 75 %MVC) across the menstrual cycle in the neutral (a) and pronated (b) forearm positions. The filled triangle denotes the mean. * Indicates the luteal phase had higher CV of force. # Indicates the follicular phase had higher CV of force.

In the neutral position at 5% (p<0.05) and 50% (p=0.05) MVC the follicular phase had greater CV of force than menses. In the neutral position at 25% (p=0.04), 50% (p=0.03) and 75% (p=0.05) MVC the luteal phase had greater CV of force compared to menses (Figure 2a). In the pronated position at 75% (p<0.001) MVC the follicular phase had greater CV of force, and at 10% (p=0.01), 50% (p<0.05) and 75% (p=0.01) MVC the luteal phase had greater CV of force compared to menses. The luteal phase at 10% (p<0.05) and 50% (p=0.04) during the pronated position had greater CV of force compared to the follicular phase (Figure 2b).

### Sex-Related Effects

There was no difference in MVC across phase in females (p>0.14) and across day in males (p>0.56) in both forearm positions (Table 1). In neutral and pronated positions, the males had a higher MVC than females (p<0.001). In males, CV of force was less in the neutral compared to pronated (p>0.05) position. Between days for males in the neutral (p<0.36) and pronated positions (p<0.72) there were no differences in CV of force. The CV of force in males from day 1 was used to assess sex-related differences in males from the three menstrual phases in females. The luteal phase in females had significantly greater CV of force in the neutral (p=0.01) and pronated (p=0.02) positions compared to males on day 1 (Figure 1).

**Table 1.**
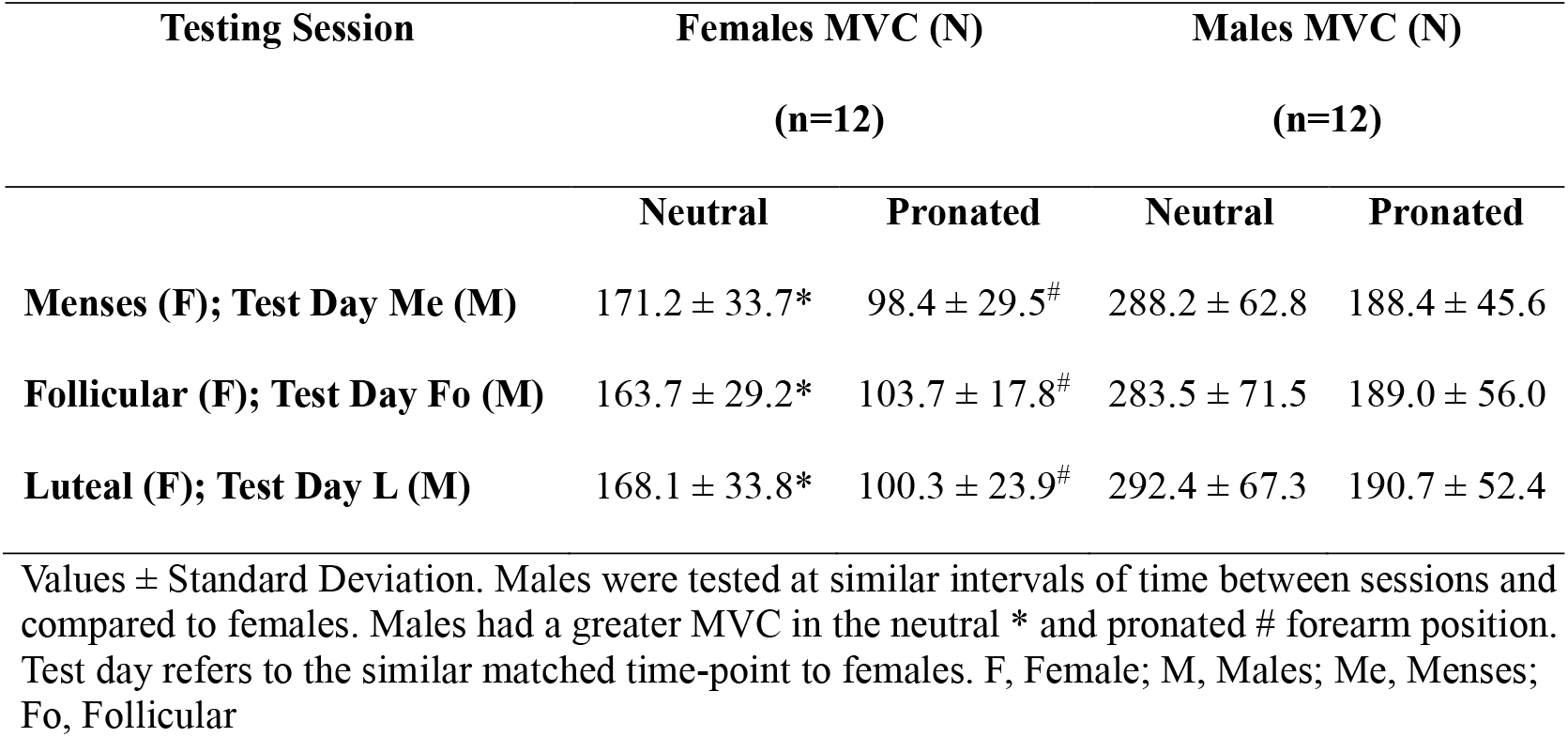
Maximal voluntary contractions across phase or test day in the neutral and pronated positions for females and males.

## Discussion

This study is the first to provide insight into the effects of menstrual phase on force steadiness in eumenorrheic females not using hormonal contraceptives, while also contributing to our understanding of sex-related differences in force control. The primary hypothesis was supported as force steadiness was lower during the luteal phase compared to menses in the neutral and pronated forearm positions. Notably, menses was the steadiest and luteal phase showed the lowest force steadiness, particularly at higher force levels. The higher force levels (i.e., 50%, 75% MVC) were steadier than low forces (i.e., 2.5%, 5%, 10%, 25% MVC) and the phase effects were most evident at greater forces. Furthermore, females in the luteal phase were less steady than males in both the pronated and neutral forearm positions. These findings underscore the importance of accounting for menstrual phase, and evaluating a broad spectrum of contraction intensities to enhance knowledge of motor controlling females.

Given that MVC is a predictor of force steadiness, it is important to consider maximal strength across the menstrual phase (Brown et al., 2010; Piasecki et al., 2023; Smart et al., 2018). Tenan et al. (2016) observed in the mid-luteal phase, when progesterone levels peak, knee extension MVC was lowest, and Rodrigues et al. (2019) reported lower strength during menses and luteal compared with the follicular phase. However, Piasecki et. al (2023) and Michalski et. al (2019) reported no change in MVC across the menstrual phase in the knee and glenohumeral extensors, respectively. Although menstrual phase effects on maximal force have been studied, findings remain inconclusive and their relevance to force steadiness is not well understood. In this study, elbow flexion MVC did not differ over the menstrual cycle or over testing sessions for males. Therefore, an increase in MVC is not the primary factor contributing to differences in force steadiness across the menstrual cycle or between males and females across the phases.

In females, sex hormone concentrations naturally fluctuate over the menstrual cycle. During the early follicular phase, estrogen is low and peaks in the late follicular phase, while progesterone remains low throughout the follicular phase (Allen et al., 2016). During the luteal phase, progesterone peaks and estrogen has a secondary lower peak relative to the follicular phase (Allen et al., 2016). Estrogen exerts an excitatory effect on the nervous system, whereas progesterone produces an inhibitory influence (Smith et al., 2002). Our results indicate that force control is optimized either when both hormone levels are low, as during menses, or when estrogen is elevated without a concurrent rise in progesterone, as in the follicular phase. Thus, progesterone could be contributing to neuronal inhibition or suppressed excitation; during the luteal phase in animal models, progesterone inhibits motor neuron excitability (Callachan et al., 1987). Inhibition, due to cyclical hormone change, could culminate in a reduction of strength and/or an increase in the variability of oscillations in common synaptic input to motor units which is associated with reduced force steadiness (Pereira et al., 2019). Force steadiness at high forces, but not low forces, is also most affected by common low frequency oscillations in motoneuron discharge rates (Dideriksen et al., 2012) and potentially contribute to the CV of force being lower in menses than the follicular and luteal phases at the highest forces of 50% and 75% MVC. To better understand if hormonal fluctuations across the menstrual cycle directly impact force steadiness, future research should investigate various force levels and common input to the motoneuron pool in relation to hormone level concentrations.

For both females and males the CV of force was greatest at the lowest force levels, gradually declined through moderate force output and then showed a slight increase at the highest force levels (see supplementary Cohen’s *d* Tables 1-8 for changes across force). The pattern of force steadiness observed in males between the two forearm positions is consistent with previous findings of isometric elbow flexion. As expected, force steadiness was less in the pronated position compared to the neutral position (Kohn et al., 2018; Smart et al., 2018; Yacyshyn et al., 2020). Importantly, force steadiness in males did not change across testing days, suggesting that the ability to maintain a steady contraction was not influenced by practice effects or repeated exposure. Therefore, the differences in force steadiness observed in females across the menstrual cycle or between sexes are unlikely to be attributed to repeat testing.

The higher CV of force in females compared to males during the luteal phase might be due to the progesterone levels. Previously, when progesterone was measured (21.4 ± 5.4 ng/dl) 10 to 3 days prior to the luteinizing hormone peak the progesterone levels did not differ from males (18.1 ± 3.1 ng/dl) (Zumoff et al., 1990a). Levels of progesterone rise substantially in the luteal phase; reaching approximately 1500 ng/dl (Zumoff et al., 1990b) contributing to reduced force control in females. Therefore, a similar level of neuronal inhibition may occur in males and females during menses and the follicular phase when progesterone levels differ minimally between the sexes. Overall, these results indicate that sex-related differences in force control is likely attributed to menstrual cycle phase, and specifically the influence of progesterone.

This study provides novel evidence that menstrual cycle phase influences force steadiness. The luteal phase, characterized by elevated progesterone, is least steady and phase effects are particularly evident at higher contraction intensities. These findings highlight the value in considering menstrual phase, and when possible, hormonal levels when evaluating sex-related differences. While males demonstrated consistent force steadiness across sessions, the variability observed in females across phases suggests a hormonally mediated modulation of motor unit behavior, potentially through changes in common inputs to motor units. Importantly, this work reinforces the value of studying females across the menstrual cycle and supports the inclusion of sex-specific and phase-specific considerations in neuromuscular research. Given this, it is imperative that future studies should directly assess hormone concentrations and neural mechanisms such as synaptic input variability to better understand how cyclical hormonal changes influence motor control across force outputs and life stages. Therefore, sex-related differences in force steadiness are likely attributed to fluctuations in peak hormone concentrations, particularly progesterone.

## Acknowledgements

The participants are thanked for the multiple visits to the lab, and timing these visits within their academic and life schedules to capture appropriate menstrual phase periods. Dr. Inglis J.G. is thanked for reading a draft of the manuscript and offering feedback.

## Notes

Research Support from the Natural Sciences and Engineering Research Council Discovery Grant Program (Grant No: 312038)

Authors disclose there are no conflicts of interest.

### Competing Interest Statement

The authors have declared no competing interest.

