## Supplemental Data 1 for "Value-Added: Importance of Incorporating Menstrual Cycle Phases to Clarify Sex-Related Differences in Force Steadiness"

**Supplementary Material**

**Supplemental Table 1.** Cohen's d values between force levels during menses in the pronated position

| Force Level | 5% MVC | 10% MVC | 25% MVC | 50% MVC | 75% MVC |
|---|---|---|---|---|---|
| 2.5% MVC | d=1.1 CI=0.43-1.67 | d=1.9 CI=1.2-2.5 | d=2.1 CI=1.3-2.8 | d=2.0 CI=1.3-2.7 | d=2.0 CI=1.3-2.8 |
| 5% MVC | | d=1.0 CI=0.4-1.6 | d=1.3 CI=0.7-2.0 | d=1.2 CI=0.6-1.8 | d=1.3 CI=0.7-1.9 |
| 10% MVC | | | d= 0.5 CI=(-0.1)-1.1 | d=0.3 CI=(-0.3)-0.9 | d=0.4 CI=(-0.2)-1.0 |
| 25% MVC | | | | d=(-0.2) CI=(-0.8)-0.4 | d=(-0.1) CI=(-0.7)-0.5 |
| 50% MVC | | | | | d=0.1 CI=(-0.5)-0.7 |

**Supplemental Table 2.** Cohen's d values between force levels during the follicular phase in the pronated position

| Force Level | 5% MVC | 10% MVC | 25% MVC | 50% MVC | 75% MVC |
|---|---|---|---|---|---|
| 2.5% MVC | d= 1.6 CI=0.9-2.3 | d=2.3 CI=1.6-3.1 | d=2.4 CI=1.6-3.1 | d=2.2 CI=1.4-2.9 | d=1.8 CI=1.1-2.5 |
| 5% MVC | | d=1.1 CI=0.5-1.7 | d=1.2 CI=0.5-1.8 | d=0.8 CI=0.2-1.4 | d=0.3 CI=(-0.3)-0.8 |
| 10% MVC | | | d=0.2 CI=(-0.4)-0.8 | d=(-0.1) CI=(-0.7)-0.4 | d=(-0.8) CI=(-1.4)-(-0.2) |
| 25% MVC | | | | d=(-0.3) CI=(-0.8)-0.3 | d=(-0.9) CI=(-1.5)-(-0.3) |
| 50% MVC | | | | | d=(-0.6) CI=(-1.2)-0.0 |

**Supplemental Table 3.** Cohen's d values between force levels during the luteal phase in the pronated position

| Force Level | 5% MVC | 10% MVC | 25% MVC | 50% MVC | 75% MVC |
|---|---|---|---|---|---|
| 2.5% MVC | d= 1.4 CI=0.7-2.0 | d=1.7 CI=1.0-2.4 | d=2.4 CI=1.6-3.2 | d=1.8 CI=1.1-2.6 | d=1.8 CI=1.1-2.5 |
| 5% MVC | | d=0.5 CI=(-0.1)-1.1 | d=1.5 CI=0.8-2.2 | d=0.7 CI=0.1-1.3 | d=0.7 CI=0.0-1.3 |
| 10% MVC | | | d=0.9 CI=0.2-1.5 | d=0.2 CI=(-0.4)-0.8 | d=0.2 CI=(-0.4)-0.8 |
| 25% MVC | | | | d=(-0.8) CI=(-1.4)-(-0.2) | d=(-0.8) CI=(-1.4)-(-0.1) |
| 50% MVC | | | | | d=0.0 CI=(-0.6)-0.6 |

**Supplemental Table 4.** Cohen's d values between force levels during menses in the neutral position

| Force Level | 5% MVC | 10% MVC | 25% MVC | 50% MVC | 75% MVC |
|---|---|---|---|---|---|
| 2.5% MVC | d= 1.9 CI=1.2-2.6 | d=2.2 CI=1.5-3.0 | d=2.7 CI=1.9-3.5 | d=2.3 CI=1.6-3.1 | d=2.3 CI=1.6-3.1 |
| 5% MVC | | d=0.5 CI=(-0.1)-1.2 | d=1.3 CI=0.7-2.0 | d=0.7 CI=0.1-1.3 | d=0.6 CI=0.0-1.2 |
| 10% MVC | | | d=0.9 CI=0.3-1.5 | d=0.2 CI=(-0.4)-0.8 | d=0.1 CI=(-0.5)-0.6 |
| 25% MVC | | | | d=(-0.6) CI=(-1.2)-0.01 | d=(-0.9) CI=(-1.6)-(-0.3) |
| 50% MVC | | | | | d=(-0.2) CI=(-0.8)-0.4 |

**Supplemental Table 5.** Cohen's d values between force levels during the follicular phase in the neutral position

| Force Level | 5% MVC | 10% MVC | 25% MVC | 50% MVC | 75% MVC |
|---|---|---|---|---|---|
| 2.5% MVC | d= 1.2 CI=0.6-1.8 | d=1.8 CI=1.1-2.4 | d=2.2 CI=1.5-3.0 | d=1.8 CI=1.1-2.5 | d=1.7 CI=1.0-2.4 |
| 5% MVC | | d=0.8 CI=0.2-1.4 | d=1.6 CI=0.9-2.3 | d=0.9 CI=0.2-1.5 | d=0.7 CI=0.1-1.3 |
| 10% MVC | | | d=0.9 CI=0.3-1.6 | d=0.1 CI=(-0.5)-0.7 | d=(-0.1) CI=(-0.7)-0.5 |
| 25% MVC | | | | d=(-0.8) CI=(-1.4)-(-0.1) | d=(-1.0) CI=(-1.6)-(-0.4) |
| 50% MVC | | | | | d=(-0.2) CI=(-0.8)-0.4 |

**Supplemental Table 6.** Cohen's d values between force levels during the luteal phase in the neutral position

| Force Level | 5% MVC | 10% MVC | 25% MVC | 50% MVC | 75% MVC |
|---|---|---|---|---|---|
| 2.5% MVC | d= 1.5 CI=0.8-2.2 | d=2.0 CI=1.3-2.8 | d=2.3 CI=1.5-3.0 | d=1.9 CI=1.2-2.7 | d=1.7 CI=1.0-2.3 |
| 5% MVC | | d=0.6 CI=0.1-1.2 | d=1.0 CI=0.4-1.7 | d=0.6 CI=(-0.02)-1.2 | d=0.2 CI=(-0.4)-0.8 |
| 10% MVC | | | d=0.5 CI=(-0.1)-1.1 | d=0.01 CI=(-0.6)-0.6 | d=(-0.3) CI=(-0.9)-0.3 |
| 25% MVC | | | | d=(-0.4) CI=(-1.0)-0.2 | d=(-0.7) CI=(-1.3)-(-0.1) |
| 50% MVC | | | | | d=(-0.3) CI=(-0.9)-0.3 |

**Supplemental Table 7.** Males Cohen's d values between force levels in the neutral position

| Force Level | 5% MVC | 10% MVC | 25% MVC | 50% MVC | 75% MVC |
|---|---|---|---|---|---|
| 2.5% MVC | d= 1.9 CI=1.2-2.6 | d=2.1 CI=1.3-2.8 | d=2.5 CI=1.7-3.3 | d=1.7 CI=1.0-2.4 | d=1.7 CI=1.0-2.4 |
| 5% MVC | | d=0.3 CI=(-0.3)-1.9 | d=1.1 CI=0.5-1.7 | d=(-0.4) CI=(-1.0)-0.2 | d=(-0.2) CI=(-0.8)-0.4 |
| 10% MVC | | | d=0.8 CI=0.2-1.4 | d=(-0.7) CI=(-1.3)-(-0.1) | d=(-0.5) CI=(-1.1)-0.1 |
| 25% MVC | | | | d=(-1.5) CI=(-2.2)-(-0.9) | d=(-1.2) CI=(-1.8)-(-0.5) |
| 50% MVC | | | | | d=0.1 CI=(-0.4)-0.7 |

**Supplemental Table 8.** Males Cohen's d values between force levels in the pronated position

| Force Level | 5% MVC | 10% MVC | 25% MVC | 50% MVC | 75% MVC |
|---|---|---|---|---|---|
| 2.5% MVC | d= 1.1 CI=0.4-1.7 | d=1.6 CI=1.0-2.3 | d-2.0 CI=1.3-2.7 | d=1.7 CI=1.0-2.4 | d=1.6 CI=0.9-2.3 |
| 5% MVC | | d=0.8 CI=0.2-1.4 | d=1.3 CI=0.7-2.0 | d=0.9 CI=0.3-1.5 | d=0.8 CI=0.2-1.4 |
| 10% MVC | | | d=0.7 CI=0.1-1.3 | d=0.1 CI=(-0.4)-0.7 | d=0.1 CI=(-0.5)-0.7 |
| 25% MVC | | | | d=(-0.6) CI=(-1.3)-0.0 | d=(-0.6) CI=(-1.2)-0.0 |
| 50% MVC | | | | | d=(-0.1) CI=(-0.7)-0.5 |
